# Characteristics of epitopes of limited variability on the head of influenza H1 haemagglutinin

**DOI:** 10.1101/2024.04.30.591841

**Authors:** José Lourenço, Hany Zinad, James Kempton, Sunetra Gupta

## Abstract

It is commonly assumed that naturally protective targets of immunity in influenza are highly variable. Theoretical work suggests, by contrast, that influenza evolution is primarily driven by naturally protective responses against epitopes of limited variability (ELV). At least one ELV has been identified on the head region of haemagglutinin of H1 influenza, opening up the possibility of producing a universal influenza vaccine. Here we demonstrate that the head region of H1 haemagglutinin can be decomposed into a number of discrete variable regions (VRs): ELVs tend to include a limited number of VRs compared to other epitopes either because of the smaller footprint of the associated antibody or because they are centred on VRs that are relatively isolated from others. We conclude that the variability of an antibody binding site is determined by the number of variable residues included in its footprint rather than the intrinsic entropy of any particular region.

## Introduction

Influenza is currently estimated to cause between a quarter to half a million deaths annually across the globe, with the majority occurring in those over 65 years of age ^1^. However, its precise impact has yet to be characterised in many parts of the world such as Africa, where much of the mortality and morbidity is concentrated in young children ^2^. The risk of complications from influenza infection also increases during pregnancy and among those with underlying medical conditions such as HIV, TB and certain immunodeficiencies ^3^.

Efforts to reduce the burden of influenza through vaccination are complicated by the ability of the virus to mutate its naturally protective targets of immunity, particularly the B-cell epitopes situated on the head of the haemagglutinin (HA) surface glycoprotein which elicit neutralising antibodies ^3^. Thus, while both inactivated and live-attenuated vaccines are made available each year, their protective efficacy can be compromised by a mismatch with the circulating strains. Vaccine effectiveness rarely exceeds 50%, falling in some years to as low as 10% ^4^, and correlations between vaccine coverage and influenza associated deaths appear to be weak ^5^. To address these issues, efforts are being made both to improve methods of production as well as to attempt to develop a “strain-transcending” universal influenza vaccine.

Current approaches towards developing a universal influenza vaccine typically rely on the possibility of artificially inducing protection against invariant targets of immunity ^4^ which are not naturally protective, such as B cell epitopes situated on the haemagglutinin stalk ^6,7^ or T cell epitopes on conserved internal antigens ^8,9^. This is because it is widely assumed that all naturally protective targets of immunity are highly variable. However, while it is necessarily true that all invariant targets of immunity must be poorly protective (as, otherwise, influenza would be like measles where a single infection would confer lifelong immunity), the converse does not holdç in other words, it is not necessarily true that all protective targets are highly variable. Indeed, we have demonstrated, through a combination of theoretical and experimental studies ^10–12^, that certain naturally protective targets of immunity located on the head of the haemagglutinin molecule are of limited variability. This opens up the possibility of making vaccines providing broad coverage without sacrificing immunogenicity ie. multivalent vaccines of durable potency with a deployable number of allelic variants.

We have identified and characterised *in vitro* and *in vivo* at least one naturally protective epitope of limited variability (OREO) of the H1 subtype of influenza A ^12^ in the head region of haemagglutinin (HA) proximal to the receptor binding site; we have performed proof-of-concept vaccine studies using the OREO epitope and are currently extending this work to identify epitopes of limited variability (ELV) in other subtypes of influenza A, as well as the Yamagata and Victoria lineages of influenza B with the aim of developing a universal vaccine against human influenza with the potential to extend to avian subtypes.

Here, we report the results of a structural bioinformatic exercise suggesting that OREO epitope itself is a mosaic of two domains of very low variability. We further demonstrate that the head region of H1 haemagglutinin can be viewed as a mosaic of variable regions (VR), and the hyper-variability observed among certain epitopes is a consequence of their footprint spanning several VRs while epitopes of limited variability (ELV) tend to contain a very low number of VRs.

## Results

### The OREO epitope in H1 haemagglutinin is a mosaic of two variable regions, VR1 and VR2

We have previously reported ^12^ that H1 haemagglutinin contains an epitope of limited variability (ELV), OREO, spanning positions 146-159 (linear numbering) in its head region, which may be grouped into five types (originally termed red, blue, green, orange and pink) on the basis of the biochemical characteristics of residues in positions 147, 156, 158 and 159.

The antigenic relationships between these variants were also interrogated through pseudotype neutralisation assays performed on mice vaccinated via a prime-boost-boost design with each variant presented on three different avian haemagglutinin scaffolds and subsequently challenged with PR8 (containing red OREO) and A/Cal/04/2009. These studies ^12^ revealed cross-reactivity between various pairs of variants, and notably also cross-protection against challenge between red and green OREO variants.

One explanation for these results is that OREO itself is a mosaic of two components which may be recognized separately or together, depending on the footprint of the antibody. OREO can be deconstructed into two variable regions, VR1 and VR2, on either end of a conserved region both in sequence space and on the structure of HA (**Figure 1**). The residues 146 and 147 together comprise VR1 which can adopt 4 subtypes, although it could be argued that the blue and green subtypes are broadly equivalent with a positive charge R/K in position 147; the red subtype contains a deletion in 147 and the pink subtype is either negatively charged or neutral. VR2 contains residues 154-158, and also can be categorised into 4 subtypes, designated as red, blue, green and orange in **Figure 1**. There appear, however, to be strong structural similarities between the orange and red variants; this is in agreement with the cross-reactivity and cross-protection observed in our previous mouse studies.

**Figure 1:**
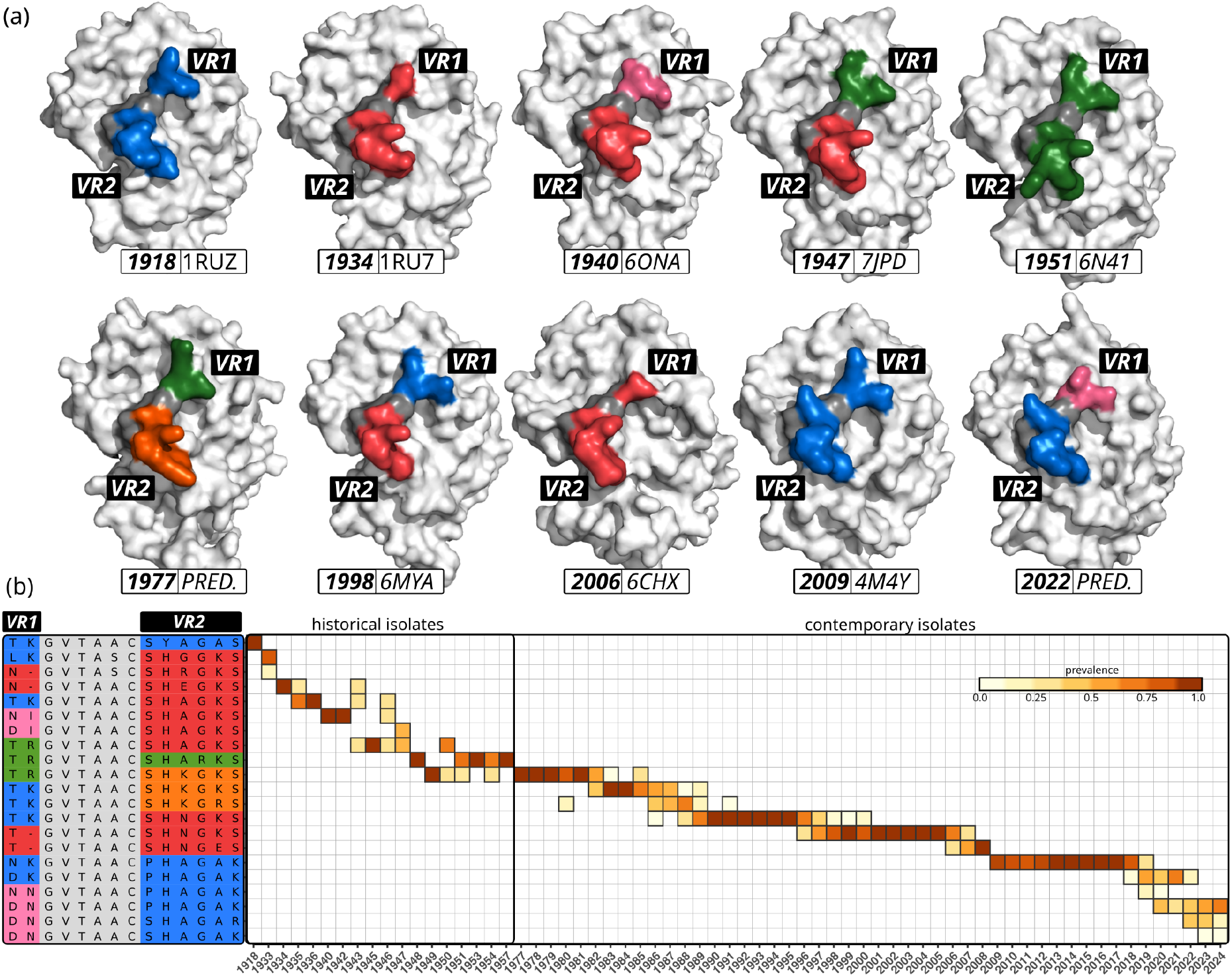
The evolution of the H1 OREO epitope from 1918 to the present: (a) The two variable regions (left side), VR1 and VR2, of OREO are tracked through time within a set of representative strains (right side): the top row contains historical strains from the period 1918 to 1957 while the bottom row contains strains from the contemporary period (since 1977). Subtypes of VR1 and VR2 are differentiated by colours (VR1: blue, red, pink, green; VR2: blue, red, green and orange) and flank an invariant region shown in grey. (b) Prevalence of different combinations of VR1 and VR2 (constituting OREO) by year among historical and contemporary isolates. Note that our reconstruction of the timeline of historical strains is based on a small number of isolates collected at irregular intervals.

The range of variation in OREO is determined by the possible combinations of VR1 and VR2. The current circulating H1 variant (since-2020) is a chimera of a pink VR1 and a blue VR2.

### VR2 constitutes a stand-alone region of diversity

We next attempted to place VR1 and VR2 in the context of the diversity exhibited across the head region of HA. We collected variable residues that were close within the AA sequence into different colour groups as shown in **Table S1**; within this scheme VR1 could be seen to belong to a larger subgroup containing variable residues 137,138,142,144,146 (shown in yellow in **Table S1**) while VR2 clearly constituted a stand-alone region of diversity spanning positions 156-159 (shown in turquoise in **Table S1**) flanked by conserved regions comprising residues 148-155 and 160-169 (163 toggles between K and R).

Figure 2. shows a range of epitopes containing VR2 as revealed by a structural bioinformatic mapping performed on PR8, using an antibody binding site diameter of 800Å. The OREO epitope, as defined in the previous section, contains both VR2 and VR1 connected by an invariant region marked in grey. Of even lower variability is an epitope we designate MAIZ, which contains only VR2, flanked by two invariant regions. Another ELV we can identify by these means is MIEZ containing VR2 and two variable residues in positions 68 and 71. By contrast, EHV1, EHV2 and EHV3 include a number of other variable residues in addition to both VR2 and VR1 which increases their potential variability. It should be noted that EHV2 maps closely to the Ca1 region within the original operational antigenic map of the HA molecule ^13,14^ obtained in the early ‘80s by comparative analysis of PR8 mutant viruses probed with monoclonal anti-HA antibodies.

### The variability of an antibody binding site is determined by the number of variable regions included in its footprint

A general principle to emerge from a comprehensive inspection of all antibody binding sites on the head of HA is that the variability of an antibody binding site is determined by the number of variable residues included in its footprint rather than the intrinsic entropy of any particular region. Epitopes of limited variability (ELVs) will thus cluster around VRs which are spatially disjoint from other VRs, such as the VR2 region (shown in turquoise in **Figures 2 & 3**) discussed in the previous section. By contrast, the variable region shown in red in **Figure 3** is in close proximity of blue and yellow VRs, as well as the purple VR region of an adjoining monomer; it is therefore difficult for an antibody to include elements of the red VR without also encompassing at least one of these other VRs. This is illustrated in **Figure 3d** by highlighting the variable residues contained within a representative selection of antibody binding sites (ABS) of diameter 800Å (each line represents an ABS with centroid in the position indicated on bold on the left).

**Figure 2:**
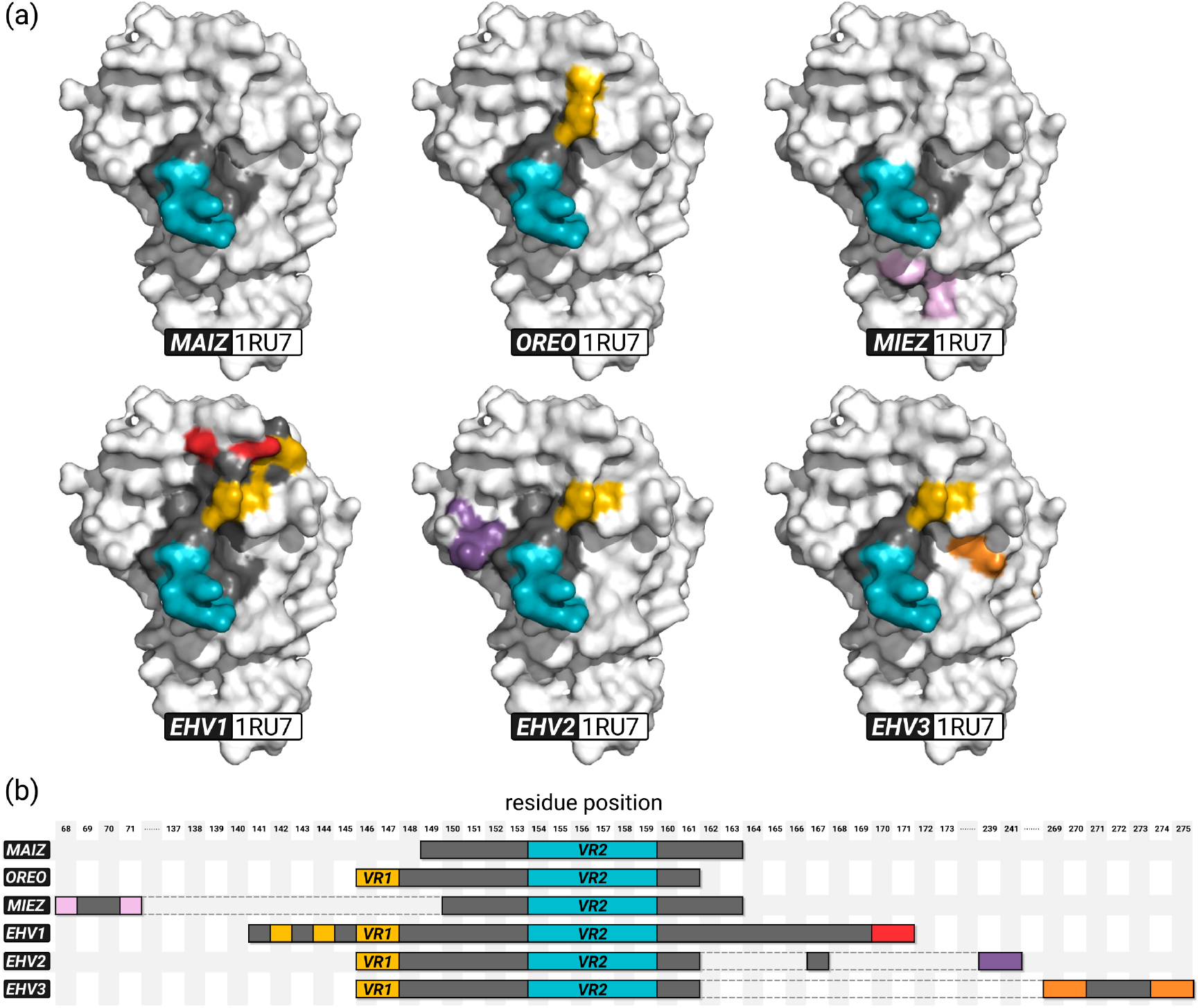
Examples of epitopes containing the VR2 region: Antibody binding sites of equal footprint size (here, 800Å and accessibility 10) positioned on VR2 can contain a variety of additional variable residues as shown here both (a) on the structure of haemagglutinin and (b) within sequence space. MAIZ, OREO and MIEZ constitute epitopes of limited variability (ELV). By contrast, EHV1-3 represent epitopes of high variability and include residues from three distinct variable regions. Colours correspond to variable residues as defined in Table S1.

**Figure 3:**
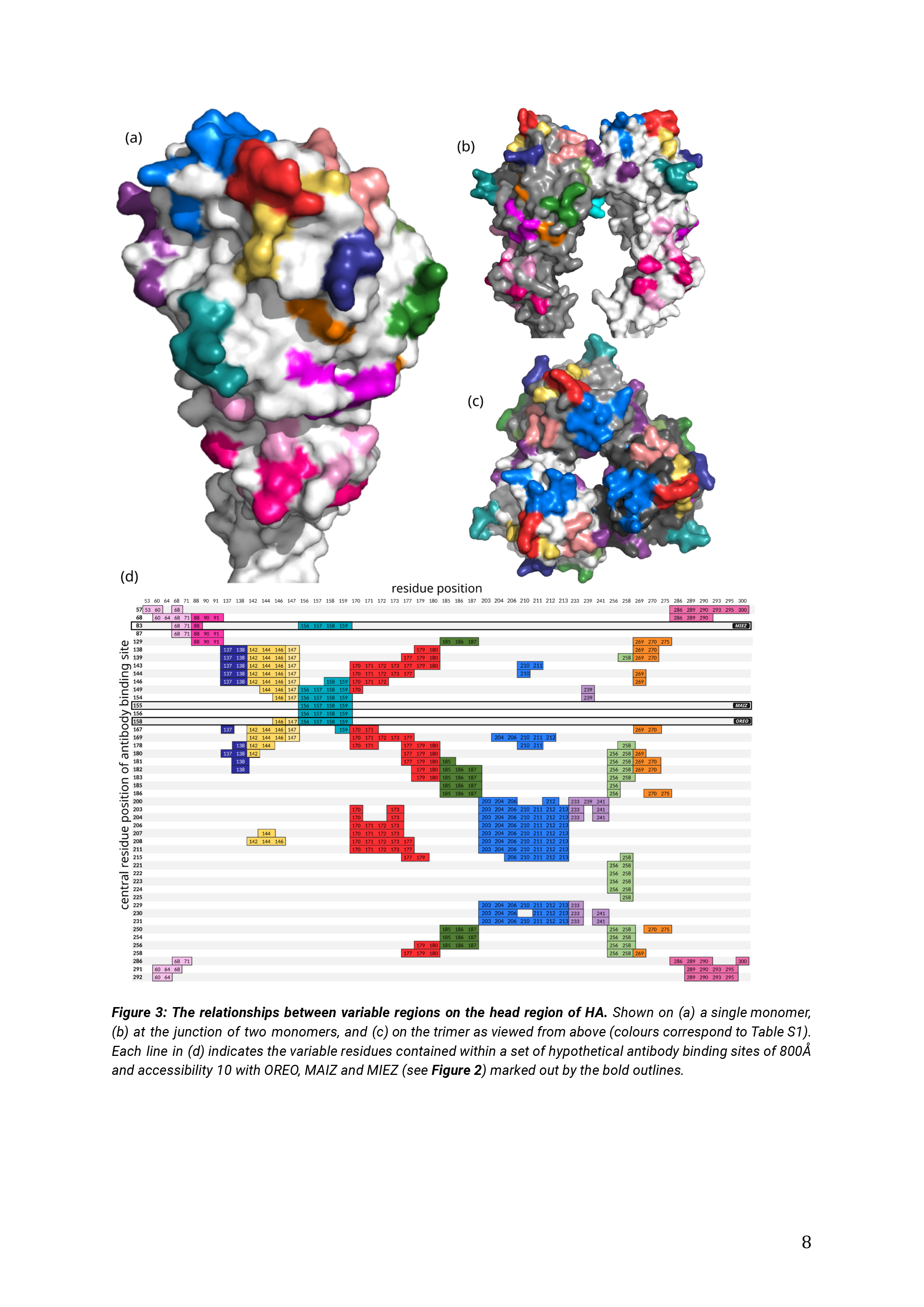
The relationships between variable regions on the head region of HA. Shown on (a) a single monomer, (b) at the junction of two monomers, and (c) on the trimer as viewed from above (colours correspond to Table S1). Each line in (d) indicates the variable residues contained within a set of hypothetical antibody binding sites of 800Å and accessibility 10 with OREO, MAIZ and MIEZ (see **Figure 2**) marked out by the bold outlines.

In line with the crystal structures shown in **Figure 3a-c**, the two-dimensional representation ABS map in **Figure 3d** locates a set of ELVs incorporating VR2 (OREO is shown for reference) but cannot identify any ELVs associated with the red VR residues as they are typically found in conjunction with blue or yellow or both, as well as with the dark and orange VRs. What distinguishes VR2 from other variable regions is that ABS centred upon it tend to remain of limited variability as the scope of potential included residues is expanded by virtue of higher ABS diameter or reduced amino acid accessibility (SASA).

## Discussion

The head region of HA can be broken down in sequence space into discrete variable regions (VR); most of these show limited variability in the combinations of constituent amino acids as may be expected under the biochemical constraints within which they have to be assembled and perform their functions (**Table S1**). Many of these VRs are, however, in close proximity, if not contiguous, within the structure of the HA molecule; in consequence, the footprint of a normal antibody is unlikely to be restricted to a single VR, thereby giving rise to a large combinatoric range of potential “epitopes”. For example, if a particular VR can only exist in 5 possible conformations and another nearby VR can exist in 4 possible conformations, then an antibody binding site that spans both will have 20 different structural variations.

How, then, do ELVs arise? One possibility is that they are the targets of antibodies of smaller footprint than the typical 700–900Å. Small footprint size may be achieved by the selective dominance of a single complementarity-determining region (CDR) loop as has been demonstrated for monoclonal antibodies such as CO5 ^15,16^ and CH65 ^17^. Both contain the VR1 portion of OREO as well as the final residue of VR2. Neutralisation patterns against H1 indicate that CO5 recognises strains with lysine in position 147 while CH65 147 is effective against strains with a deletion in position 147 and unable to neutralise strains with either a lysine or arginine in this position; these results are in agreement with the analysis of the OREO epitope presented in **Figure 1** and are supported by neutralisation patterns of other “broadly neutralising” monoclonal antibodies such as 5J8 ^18^ and IFI which also include position 147 ^19^.

These monoclonal antibodies exhibit extensive (even heterosubtypic) cross-neutralising properties on account of including only a limited number of residues surrounding the receptor binding site (**Table S1**)). As such, they constitute ELVs rather than bNabs by virtue of their smaller footprints. By contrast, the antibody binding sites pivoting upon VR2 (**Figure 2**) tend to be of limited variability, even when their footprint is large, owing to the relative isolation of VR2 which is flanked by conserved regions within the structure of HA, even though it sits upon the lip of the crater which contains the receptor binding site (RBS). Antibody binding sites centred on VR2 may extend to contain some elements of other VRs and yet still retain low overall variability, as exemplified by the epitope OREO which we have identified as a potential vaccine candidate ^12^.

Of the classical antibody binding sites ^12–14,20^, only Cb is composed of a single VR (light pink in **Figure 3** and **Table S1**) which can be resolved into 5 distinct variants; however, we could not find any ABS in our structural bioinformatic analysis (**Figure 3**) which included only Cb on account of its close proximity to the other pink VRs. The sites Sa, Sb and Ca (as defined in ^20^) include more than one VR and therefore exhibit higher variability, although with clear evidence of periodic re-emergence of the same variants. An extensive epitope mapping exercise using escape mutants by Matsuzaki et al ^21^ confirms that the bulk of antibodies are restricted to the Sa, Sb and Ca2 (a part of Ca that includes VR2) but they were also able to identify at least one antibody which included Ca2 and position 147 (corresponding to OREO) and also a “broadly neutralising” monoclonal antibody (n2) centred on the RBS similar to those described above ^15–19^. By contrast, the monoclonal antibody 2D1, derived from B cells of survivors of the 1918 H1N1 pandemic ^20^, is highly diverse on account of possessing a footprint that spans 5 VRs; consequently it has low affinity for all strains except 2009 H1 HA (CA04) which is identical in every single residue.

Our analysis challenges the conventional wisdom that low variability is an indication of low natural potency (**Figure 4**). Instead, we show that epitopes of low variability can be equally as protective as those of high variability, and therefore under equally high selection pressure. Our framework explains such puzzling observations as “very little drift occurred in the Ca site despite the relatively high virus-neutralizing of anti-Ca antibodies *in vitro*” ^13^, by demonstrating that it is the architecture of that region that determines its variability rather than level of selection pressure. Indeed, it is a common misconception that the absence of variability indicates that a site is under lower immune selection; they are the structural constraints that primarily dictate whether immune evasion is possible (cf. for the measles virus, viral escape from neutralization is impossible as it leads to loss of receptor-binding activity ^22^).

**Figure 4:**
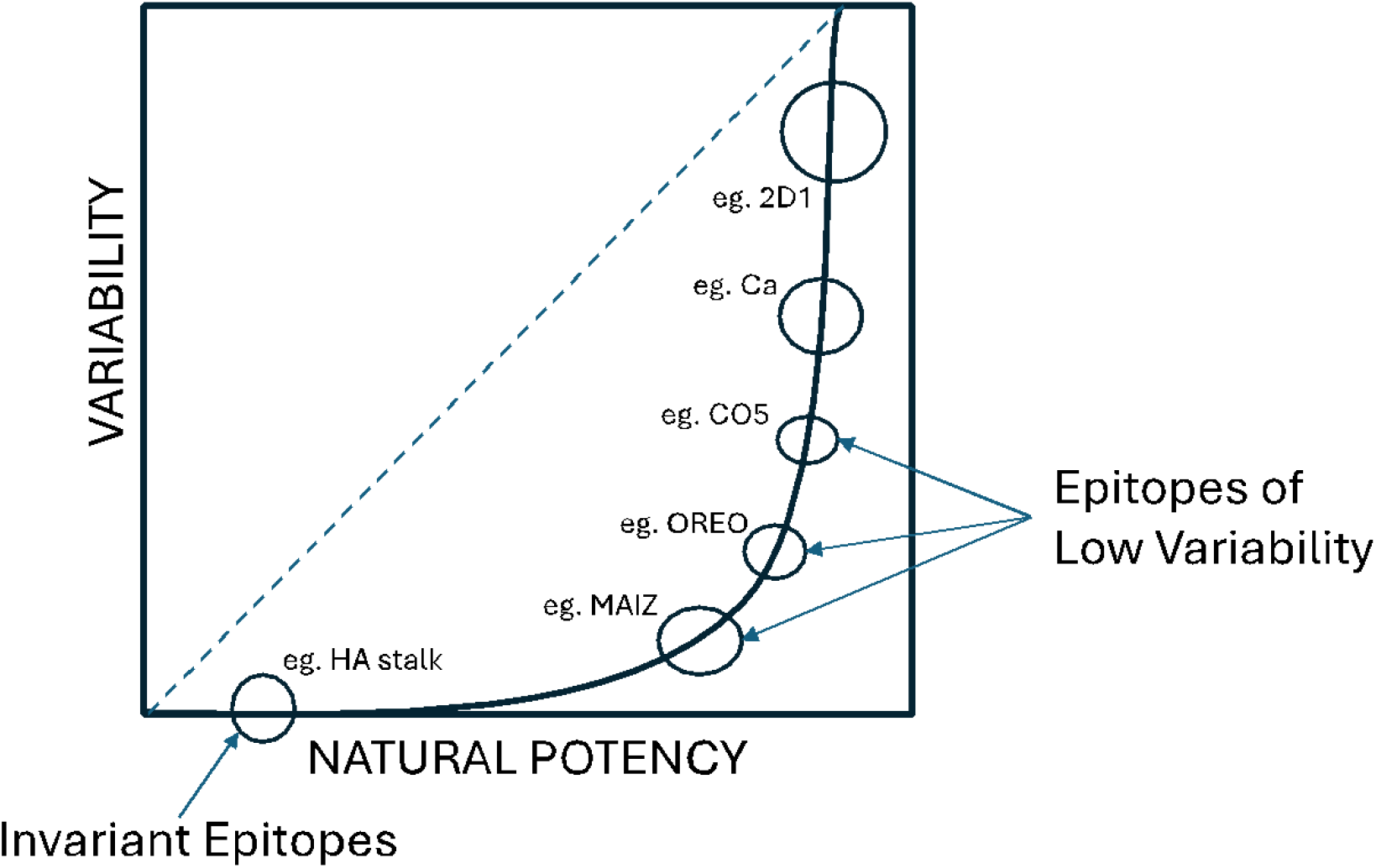
Schematic representation of the relationship between epitope variability and potency within our framework. The dashed line represents a scenario where potency increases linearly with variability.

An ELV under strong immune selection will cycle through its various conformations, as shown in **Figure 1**. This is manifest in patterns of cross-neutralisation as observed by us ^12^ between A/Solomon Islands/2006, PR8 and A/WSN/1933 which all share red OREO, as well as cross-protection in mice from lethal PR8 challenge upon vaccination with the 2006 red OREO variant. Vaccinating mice with a 1918-like strain has been shown to confer protection against lethal challenge with the 2009 H1N1 strain of comparable levels to homologous inoculation ^23^; both 2009 and 1918 contain the blue OREO variant, as do the two swine viruses (Sw/30 and NJ/30) used in the same experiment which also gave high levels of protection. A range of other historically distributed human viruses (Wei/43, USSR/77, TX/91, Bris/07) which do not contain blue OREO showed more rapid weight loss and slower recovery, indicating that (i) antibodies against OREO play an important role in protection and (ii) there are other shared ELVs (as predicted by the model ^11^ and other experimental studies ^11,24^) which also contribute to protection. The re-emergence of blue OREO in the 2009 pandemic strain is also suggested by the high prevalence of neutralising antibodies to it in older individuals who were likely to have been naturally exposed to the 1918 strain ^25,26^. The newly reported H1N2 strain from November 2023 ^27^ contains an OREO epitope that is effectively identical to the variant circulating between 1989-1996 and also between 1935-42 (**Figure S1**).

It has recently been shown ^28^ that sequential vaccination with a set of H1N1 strains is able to induce a wide spectrum of neutralising antibodies and protection against challenge with heterologous strains in swine. An inspection of the HA sequences used in this study reveals that all VR2 variants identified in the human H1N1 data were covered, thereby suggesting that at least part of the protection derived from neutralising antibodies targeting an ELV containing VR2. Sequential infection of ferrets, both with historical and contemporary H1N1 strains has also been shown to confer some protection against weight loss upon challenge with 2009 H1N1, as well as to significantly reduce ability to transmit to naïve ferrets ^29^; the sera of ferrets infected with multiple strains shown to induce cross-reactive antibodies to 2009 H1N1 however as none of the “vaccine” strains shared the same VR2 region as 2009 H1, this must have derived from antibodies against other ELVs. These experiments suggest that a straightforward option for exploiting the existence of ELVs is to vaccinate individuals with a cocktail of relevant strains. However, we believe that using a prime-boost strategy to focus the response on a particular ELV ^12^ is more likely to yield robust and durable protection.

In summary, our methodology opens up a novel pathway to delivering universal protection against all strains of influenza through intelligent vaccine design. An additional advantage of this approach is that it exploits the potential of natural immunity rather than artificially stimulating responses that are not naturally protective. Alternative approaches that computationally engineer artificial vaccine constructs ^30,31^ may elicit broadly cross-reactive antibodies and may eventually provide universal coverage against influenza, but we believe that it will be far more effective to achieve broad and durable protection by targeting naturally protective epitopes of limited variability on the head of the influenza haemagglutinin surface antigen.

## Methods

### ABS analysis

This was achieved by mapping hypothetical antibody-binding sites onto the crystal structures of H1 HA and calculating the amino acid sequence variability contained within each site with reference to a set of available sequences. Analysis included the entire head from C59 to C292 for H1N1 viruses ^32^.

### Obtaining structures

Where available, crystal structures were used for analysis. In some cases, structures deposited in the PDB contained truncated amino acids and these were replaced with complete residues in COOT (Crystallographic Object-Oriented Toolkit). Where crystal structures were not available, AlphaFold2 models were used.

### Structural bioinformatic analysis

For the structural bioinformatic analysis we used the publicly available structures (at www.rcsb.org/): 1RUZ (A/South Carolina/1/1918(H1N1)), chains H, J, L), IRU7 (A/Puerto Rico/8/1934(H1N1), chains A, C, E, G, I, K), , 6CF7 (A/Solomon Islands/3/2006(H1N1), chain A) and 3LZG (A/California/04/2009(H1N1), chains A, C, E, G, I, K ). The analysis followed a pipeline with X steps: (1) quantifying accessibility of amino acid residues (AAR), (2) defining antibody binding sites (ABS), (3) aggregating results across structures and their chains. For step (1), the accessibility of AAR on H1N1 HA (for all structures and chains independently) was quantified using the Solvent Accessible Surface Area (SASA) C library (v2.0.3, www.freesasa.github.io). In step (2), for every AAR on H1N1 HA (for all structures and chains independently) an spherical area centred at the AAR (ABS pin) with specific radius (in Å) was considered and all AAR within the area were identified (using as references the positions of each AAR’s alpha-carbon). Finally, in step (3), for each combination of considered AAR accessibilities (1, 10, 30) and spherical radiuses (500, 800, 1000) (**Figure S2**), AARs belonging to ABSs were identified if present in at least one chain of any structure (**Figure S3**).

## Acknowledgements

We are grateful to Matt Higgins for his help with the structural analysis and to Paul Klenerman and Matt Edmans for their support of the project. The authors received financial support from Blue Water Vaccines.

## Author contributions

Conceptualization: JL, SG. Methodology: JL, SG, AZ. Software: JL, SG. Validation: JL, SG, AZ, JK. Formal Analysis: JL, SG. Investigation: JL, SG, AZ. Resources: SG. Data Curation: JL, SG, AZ, JK. Writing – Original Draft: JL, SG. Writing – Review & Editing: JL, SG, AZ, JK. Visualization: JL, SG. Supervision: SG. Project Administration: SG.

## Supplementary material

**Supplementary Figures** (word document): including Supplementary Figures S1-3.

**Supplementary Tables** (excel document): including Supplementary Table S1 - Variable residues that were proximal within the AA sequence were collected into different colour groups. Classical binding sites Sa, Sb, Ca, Cb (as defined in (Xu et al. 2010)) are shown across the top, and the binding sites of monoclonal antibodies discussed in the text are shown below the sequences, as is OREO. Grey boxes correspond to conserved residues included in the footprint of an antibody.

