## Supplementary Figures for "Characteristics of epitopes of limited variability on the head of influenza H1 haemagglutinin"

### Supplementary Figures S1-3

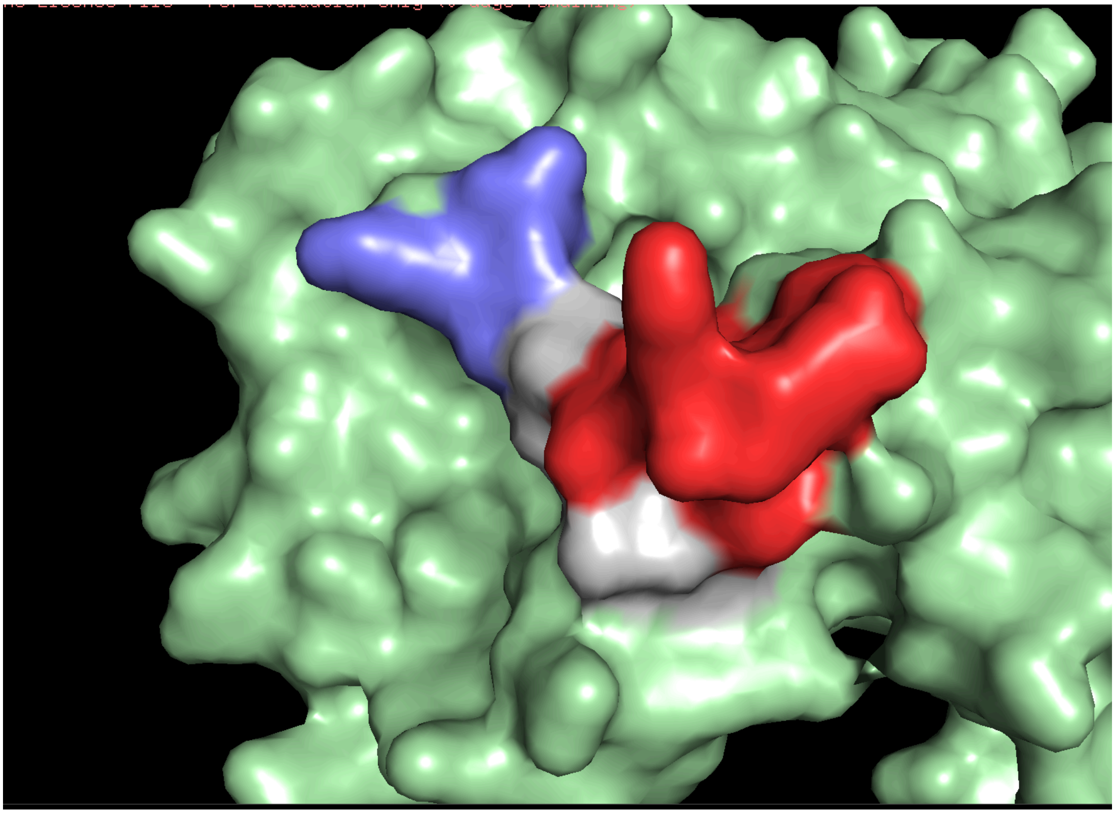

**Figure S1**: The newly reported H1N2 strain from November 2023 (Cogdale et al. 2024) contains an OREO epitope that is effectively identical to the variant circulating between 1989-1996 and also between 1935-42.

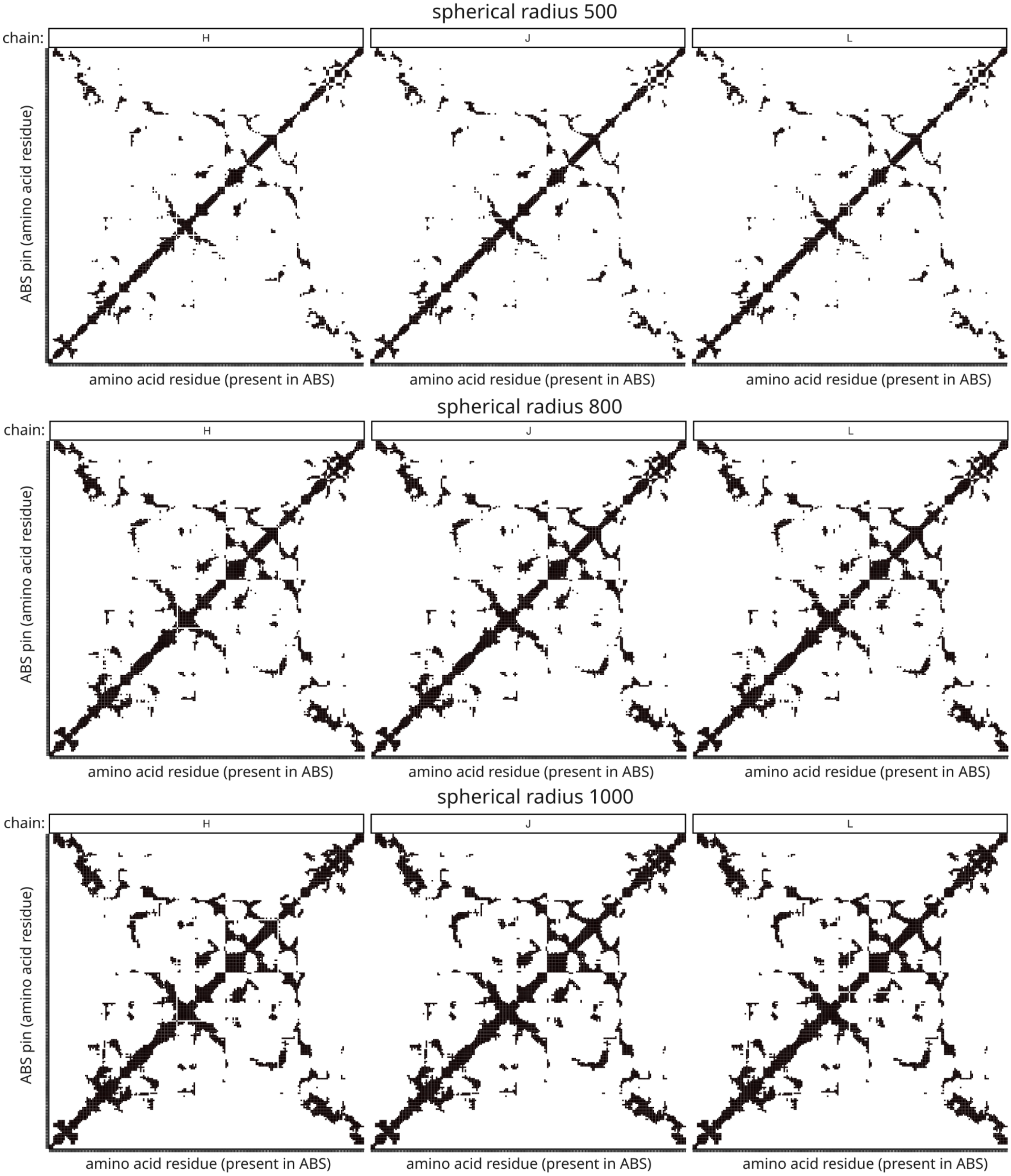


**Figure S2:** For every amino acid residue (AAR) on H1N1 HA an spherical area centred at the AAR (y-axis, the antibody binding site, ABS, pin) with specific radius (in Å) was considered and all AAR within the area were identified (using as references the positions of each AAR's alpha-carbon). Cells (X-Y positions) in black mark AARs present in the corresponding ABS. This figure includes as an example, the results from structure 1RUZ and its H, J and L chains.

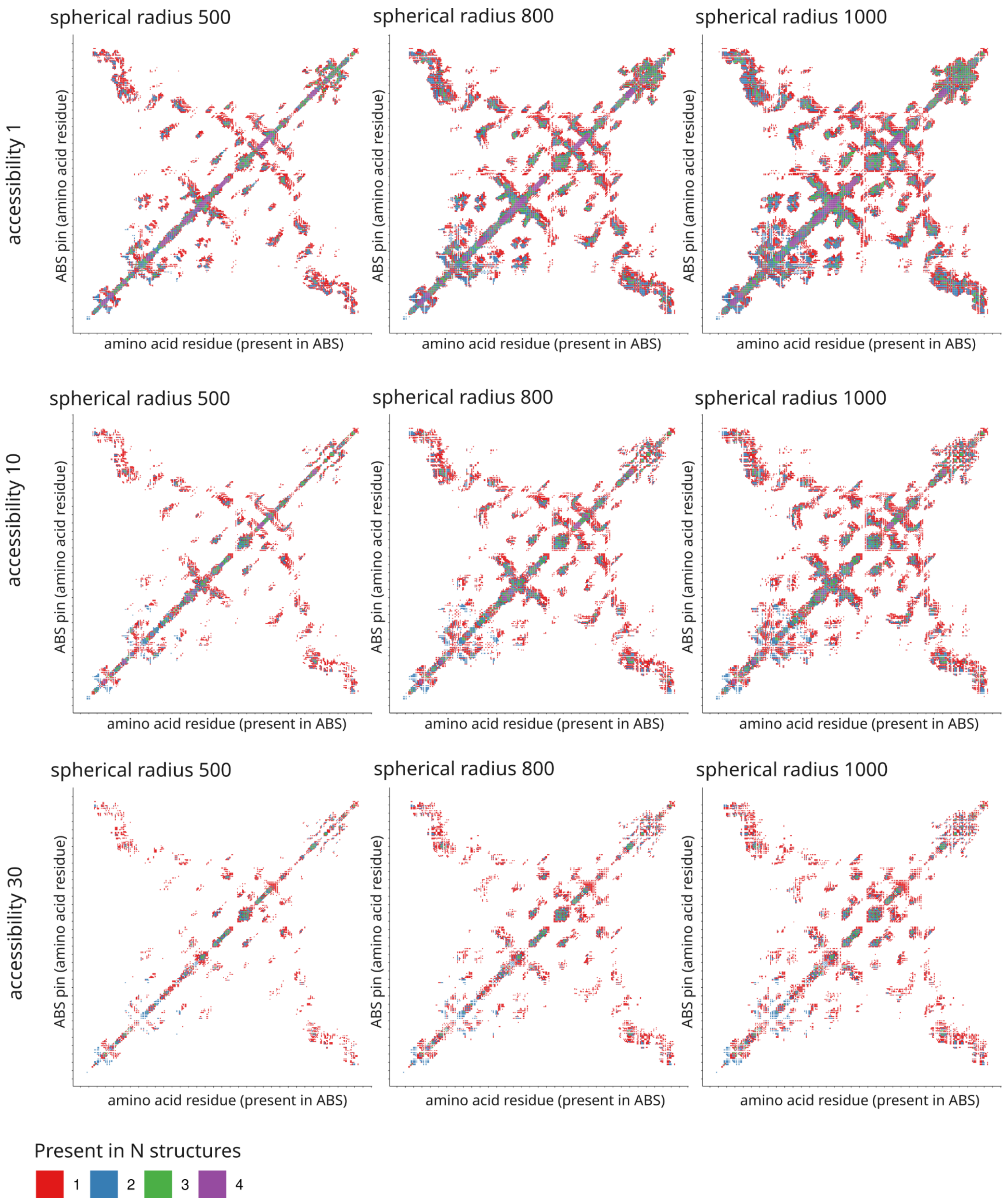


**Figure S3:** For each combination of considered amino acid residue (AAR) accessibilities (1, 10, 30) and spherical radiuses (500, 800, 1000), AARs belonging to antibody binding sites (ABS) were identified if present in at least one chain of any structure. Cells (X-Y positions) are here coloured according to whether they were present in 1-4 structures.
